# Using The Cancer Genome Atlas from cBioPortal to Develop Genomic Datasets for Machine Learning Assisted Cancer Treatment

**DOI:** 10.1101/2025.02.17.638660

**Authors:** Abu Asaduzzaman, Christian C. Thompson, Fadi N. Sibai, Md J. Uddin

**Affiliations:** Department of Electrical and Computer Engineering, Wichita State University, Wichita, Kansas 67260, USA; College of Engineering and Architecture, Gulf University for Science and Technology, West Mishref, Hawally 32093, Kuwait; Department of Biochemistry, Vanderbilt University School of Medicine, Nashville, Tennessee 37232, USA

**Keywords:** genomic dataset, PolyPhen, SIFT, random forest (RF), extreme gradient boosting (XGBoost), ensemble learning

## Abstract

Predicting the impact of genetic mutations is crucial for understanding diseases like cancer. Polymorphism Phenotyping (PolyPhen) and Sorting Intolerant From Tolerant (SIFT) are key tools for assessing how amino acid substitutions affect protein function and mutation pathogenicity. To our knowledge, no ready-to-use genomic dataset exists for prediction models to identify potentially harmful mutations, which could support research and clinical decisions. This study develops genomic and non-genomic datasets using The Cancer Genome Atlas (TCGA) from cBioPortal and applies machine learning models to predict PolyPhen and SIFT scores. We explore three classification models: Random Forest (RF), Extreme Gradient Boosting (XGBoost), and an ensemble RF-XGBoost model. Experimental results show that genomic data yields more accurate predictions than non-genomic data. The ensemble RF-XGBoost model performs best on genomic data, achieving average accuracies of 88.43% for PolyPhen and 95.13% for SIFT, highlighting the potential of artificial intelligence in genetic mutation analysis for disease treatment.

## 1. INTRODUCTION

Cutaneous melanoma, a type of skin cancer, originates from the pigment-producing cells called melanocytes. It is generally caused by ultraviolet (UV) radiation from natural sunlight and indoor tanning; however, there are additional contributing factors that are unrelated to ultraviolet radiation exposure such as genetic predisposition, family history of melanoma, hormonal changes, and aging (9, 31, 32). Recent advancements in genomics have aided in the identification of mutations associated with skin cutaneous melanoma, offering useful insights into potential therapeutic targets and prognostic indicators. **Figure 1** shows a conceptual illustration of the pathogenesis and genomic insights of cutaneous melanoma. PolyPhen and SIFT scores are important tools that assess the functional impact of mutations on protein structure and function (36, 40).

**Figure 1.**
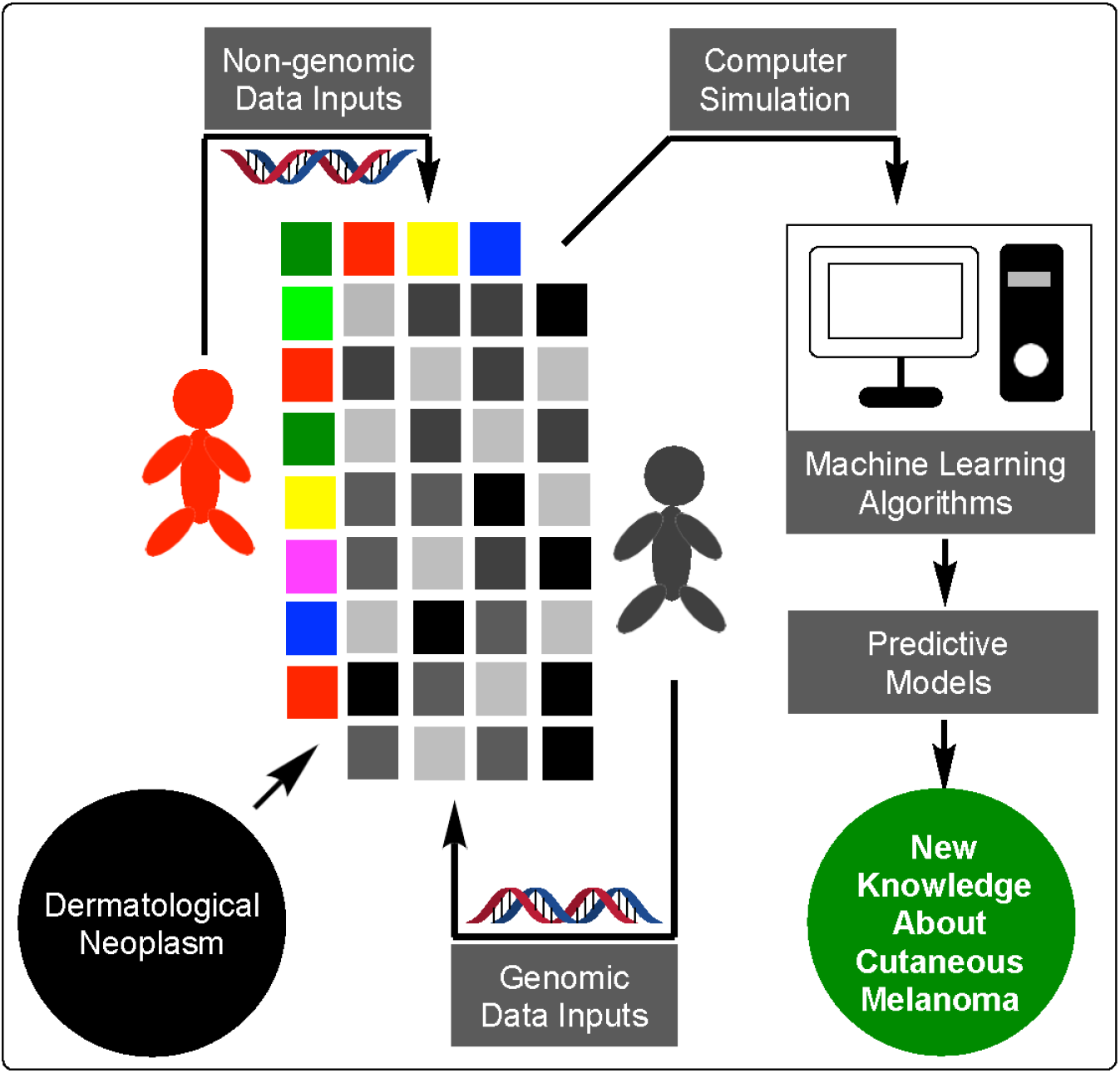
Overview of cutaneous melanoma development, including genetic mutations and functional mutation assessment.

Studies show that the growing demand for time-efficient error-free solutions to cancer diagnosis and treatment is fueling the interest in developing machine learning models to accurately classify cancerous cells (19, 59). Machine learning techniques have emerged as powerful tools in genomic research, enabling prediction of mutation effects with increased accuracy. Recent studies highlight the significant potential of genomic data to enhance cancer detection, diagnosis, and treatment (17, 28, 57). However, challenges remain, particularly regarding the quality of data and the ability of models to accurately predict the functional impact of mutations, which must be addressed to fully realize the potential of genomic datasets.

This study aims to develop effective genomic datasets for machine learning applications using data from TCGA, which provides interactive exploration of multidimensional cancer datasets molecularly characterized across various cancer types. The effectiveness of these genomic datasets in predicting PolyPhen and SIFT scores for skin cutaneous melanoma mutations is evaluated using machine learning models. We investigate which dataset—genomic or non-genomic—provides the best predictions for PolyPhen and SIFT scores using three popular machine learning algorithms: RF, XGBoost, and an ensemble RF-XGBoost model. Extensive data preprocessing and correlation analysis are employed to ensure the integrity of both genomic and non-genomic datasets. Additionally, the datasets are normalized to mitigate the risk of model overfitting.

The organization of this paper is as follows: Section two provides a review of the background materials. Section three presents the approach to assess the impact of genomic datasets for predicting the PolyPhen and SIFT scores. The experimental setup and results are discussed in Sections four and five, respectively. Finally, Section six concludes the paper.

## 2. BACKGROUND MATERIALS

In this section, we review existing literature on genetic mutations in cancer, the TCGA dataset, PolyPhen, SIFT, and machine learning techniques to provide context for the proposed approach.

### 2.1. Genetic Mutations in Cancer

Genetic mutations are alterations in the deoxyribonucleic acid (DNA) sequence of a gene. In the context of cancer, these mutations can disrupt normal cell functions, leading to uncontrolled growth and tumor formation. Skin cutaneous melanoma is a type of cancer that begins in melanocytes, the cells responsible for producing melanin which gives the skin its color (30, 31, 39, 48). It is one of the most aggressive and deadly forms of skin cancer, primarily caused by ultraviolet (UV) radiation exposure from natural sunlight and indoor tanning. Although skin cutaneous melanoma comprises less than 5% of malignant skin tumors, it is responsible for about 60% of lethal skin cancers (33, 48). Early detection and treatment are crucial, as a progressed melanoma can be challenging to treat and often has a poor prognosis.

The pathogenesis of melanoma is complex, involving both genetic and environmental factors. Genetic mutations play a critical role in the initiation and progression of melanoma. These mutations can occur in various genes that regulate cell proliferation, apoptosis, and deoxyribonucleic acid (DNA) repair. Several studies have shown that melanoma spread results from genetic mutations and alternations in the tumor microenvironment, characterized by the overexpression of proteins that promote tumor invasion and infiltration of surrounding tissues (12, 15, 37, 50). These changes can lead to loss of function, gain of function, or dominant-negative effects, contributing to cancer progression (18, 49, 52). In our work, we assess the effects of genetic mutations in skin cancer by using both genomic and non-genomic datasets in classification models.

### 2.2. The Cancer Genome Atlas (TCGA) Dataset

Cancer is essentially a genomic disease, caused by mutations in a cell’s DNA, ribonucleic acid (RNA), and proteins that cause cells to grow excessively. In this study, we utilize the skin cutaneous melanoma data from the TCGA dataset (55, 58). The Pan-Cancer Atlas project within TCGA, accessible through the cBioPortal website, comprises approximately 8,942 tumors spanning 32 different cancer types. Collectively, these tumors exhibit around 5,717,732 somatic mutations. The comprehensive dataset provides a wealth of genomic and non-genomic information, enabling detailed analysis of the genetic mutations of skin cutaneous melanoma. The integration of this dataset into our study facilitates a deeper understanding of the disease, supporting the development of more precise models and personalized treatment strategies.

### 2.3. Polymorphism Phenotyping (PolyPhen)

PolyPhen is a tool designed to predict the potential effects of amino acid substitutions on the structure and function of human proteins. It helps in assessing whether a particular genetic mutation could be benign, possibly damaging, or probably damaging (1, 2, 16). In modern human genetic research, PolyPhen-2 is developed to predict the effects of amino acid substitutions using a high-quality multiple protein sequence alignment pipeline and a prediction method based on a supervised machine-learning classification algorithm, specifically the Naïve Bayes Classifier (2, 5, 41). The PolyPhen-2 score ranges from 0.0 (tolerated) to 1.0 (deleterious). Variants with scores of 0.0 are predicted to be benign, while values closer to 1.0 are more confidently predicted to be deleterious. The score can be interpreted as follows: Scores from 0.0 to 0.15 indicate that the variants are predicted to be benign. Scores from 0.15 to 0.85 suggest that the variants are possibly damaging. Scores from 0.85 to 1.00 indicate that the variants are more confidently predicted to be probably damaging (1, 16). The TGCA dataset provides PolyPhen values categorized as benign, possibly damaging, probably damaging, and unknown. In our work, we exclude the datapoints associated with the unknown category.

### 2.4. Sorting Intolerant from Tolerant (SIFT)

SIFT predicts the functional impact of amino acid substitutions in proteins by analyzing sequence homology and evaluating the physico-chemical similarities between the original and alternate amino acids (42, 43, 54). SIFT scores range from 0.0 to 1.0, with lower scores indicating a higher likelihood of the amino acid substitution being damaging. Scores from

0.00 to 0.05 are predicted to be damaging and likely to affect protein function, as these changes occur at positions that are highly conserved and critical for the protein’s structure or function. Scores from 0.05 to 1.00 are considered tolerated and are less likely to affect protein function, as these positions are less conserved and can accommodate a variety of amino acid changes without significant impact (1, 22, 42, 51). The TGCA dataset contains four qualitative prediction values: Deleterious, Deleterious - low confidence, Tolerated, and Tolerated - low confidence. In our work, we exclude the low confidence classes, as these labels indicate a lower level of certainty due to limited sequence data or variable conservation.

### 2.5. Machine Learning in Genomic Research

In this study, we develop genomic and non-genomic datasets to train machine learning models for predicting PolyPhen and SIFT scores associated with cancer mutations and assess the effectiveness of the genomic datasets. Three popular machine learning classification models—RF, XGBoost, and an ensemble RF-XGBoost model—are employed to predict the PolyPhen and SIFT scores using both genomic and non-genomic data.

#### 2.5.1. Random Forest (RF)

RF is a powerful and versatile classification model known for its ability to handle missing data effectively (20, 25, 46). By constructing multiple decision trees and averaging their predictions, RF mitigates the impact of incomplete data, making it particularly suitable for datasets with missing values, such as the TCGA dataset. This inherent flexibility allows RF to maintain performance even when certain data points are absent, as it can continue to build accurate predictions by leveraging the remaining features. However, due to the model’s effectiveness on smaller datasets, it is essential to be cautious in how we approach data preprocessing. Over-aggressive data cleaning techniques could result in a loss of valuable information, which may negatively affect the RF model’s predictive performance. Careful consideration must be given to the balance between eliminating noise and preserving crucial details within the dataset, as this balance is critical to maximizing the accuracy and reliability of the model. Proper preprocessing ensures that the model has sufficient and relevant data to extract meaningful patterns, especially when working with limited datasets. RF offers an effective approach to predicting PolyPhen and SIFT classifications, providing robust performance, resistance to overfitting, and scalability.

#### 2.5.2. Extreme Gradient Boosting (XGBoost)

XGBoost is a highly effective model known for its exceptional ability to handle imbalanced datasets, making it particularly valuable in scenarios where class distributions are disproportionate (6, 11). This capability will be especially helpful in the TCGA data, where the PolyPhen and SIFT classes exhibit varying distributions. The model’s robustness ensures that it can maintain accuracy across these classes, effectively addressing the challenges posed by minority class representation during the training process. However, XGBoost is also sensitive to noisy data, which means we must ensure that the TCGA data is properly preprocessed to remove irrelevant or misleading features. Effective data cleaning and feature selection are crucial to mitigate the risk of overfitting, as the model may otherwise latch onto noise instead of meaningful patterns. By carefully curating the dataset, we can enhance the model’s performance and ensure that it capitalizes on the relevant information available, ultimately leading to more accurate predictions for PolyPhen and SIFT classifications. XGBoost provides a powerful method for predicting PolyPhen and SIFT classifications, demonstrating high accuracy, exceptional handling of imbalanced datasets, and resilience against overfitting. Its efficient algorithms enable rapid training and deployment, making it an ideal choice for complex genomic data analysis.

#### 2.5.3. Ensemble Model of RF and XGBoost

An ensemble model that combines RF and XGBoost effectively leverages the strengths of both models, enhancing predictive accuracy and robustness for PolyPhen and SIFT classifications (10, 34, 44). By harnessing the unique advantages of RF’s ability to manage missing data and XGBoost’s proficiency in handling imbalanced datasets, the ensemble model can achieve superior classification accuracy, often surpassing the performance of each model when used individually. Moreover, the ensemble approach improves robustness against overfitting by capitalizing on the inherent differences in the learning processes of the two models. Each model may capture different aspects of the data, leading to a more comprehensive understanding of the underlying patterns. In this context, blending—a technique that combines the predictions of multiple models by averaging their outputs—allows for a more nuanced final prediction that leverages the strengths of each model (7, 24, 35, 45).

However, implementing ensemble models necessitates rigorous tuning and careful integration, which can be resource intensive. The computational resources required for training are significant, and ensuring compatibility and coherence between the RF and XGBoost components is critical for optimal performance. Striking the right balance between their contributions can also present challenges, requiring meticulous adjustments to hyperparameters and thorough validation. Nevertheless, with appropriate optimization strategies and validation techniques in place, an ensemble of RF and XGBoost can serve as a highly effective tool for predicting PolyPhen and SIFT classifications.

## 3. PROPOSED APPROACH FOR CONSTRUCTING AND EVALUATING GENOMIC DATASETS

In this section, we describe the proposed approach, which begins with multiple data files containing both genomic and non-genomic data from various medical tests. **Figure 2** illustrates the primary steps of the proposed approach, including the integration of data points from various files into a single file using PATIENT ID; data preprocessing; splitting the data into genomic and non-genomic types; feature selection; correlation analysis; resampling to address data imbalance; and applying classification models. It should be noted that the split data can be used for classification models without feature selection, correlation analysis, and/or resampling. Once predictions due to both genomic and non-genomic datasets meet acceptable thresholds, the results are analyzed to evaluate the impact of the genomic dataset on predicting the PolyPhen and SIFT scores.

**Figure 2.**
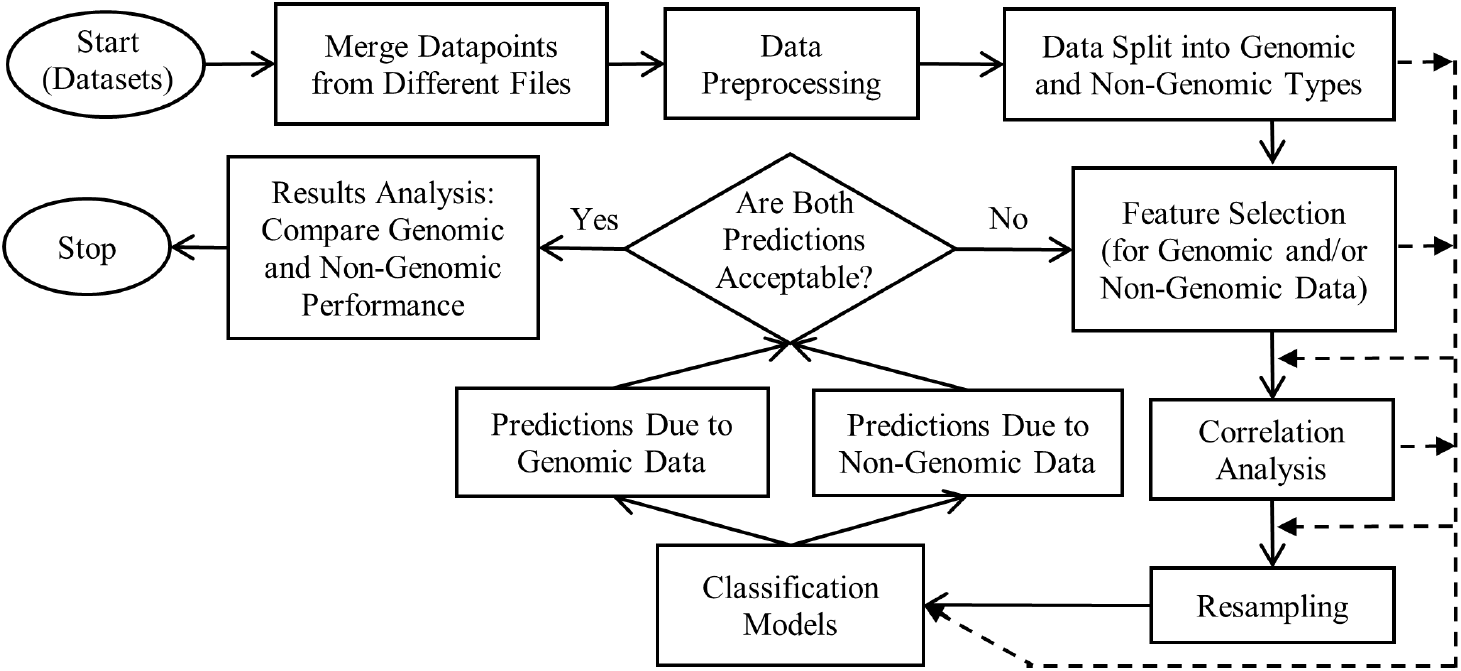
Major Steps of the Proposed Approach to Assess the Impact of Genomic Data

### 3.1. Merge Datapoints from Different Files

In this step, we study the TCGA datasets from cBioPortal and select 47 data files, which include both license and text files. Closer inspection confirms that most of the files contain limited or irrelevant information, which makes them unsuitable for combining into a cohesive data set for our research. After conducting a thorough review and analysis of the data files, we identify five files that would contribute significantly to this study. These files include data clinical patient.txt that provides essential clinical information on the patients and data mutations.txt that captures genetic mutation data. File data clinical sample.txt contains relevant clinical sample information and data clinical supp hypoxia.txt offers supplementary clinical data, specifically focusing on hypoxia conditions. Finally, data timeline sample acquisition.txt provides detailed timeline data on sample acquisition. These five files are deemed essential for constructing a master dataset to be used in machine learning analysis.

### 3.2. Data Preprocessing

In this crucial step, the raw data is converted into a clean, structured format suitable for machine learning models. First, in data cleaning, missing values are addressed to enhance the dataset’s integrity. All empty rows in the PolyPhen and SIFT columns are removed, resulting in a reduction of the dataset rows. This step is vital because retaining empty rows could lead to inaccuracies in model predictions. To further refine the dataset, columns with a large number of missing values are dropped. Eliminating these columns is essential to avoid introducing noise that could distort model training. Additionally, missing values in string columns are replaced with ‘No data available’ or ‘Unknown’ to maintain consistency, which helps to ensure that the dataset remains interpretable and usable. Columns that contained only one type of information are also eliminated, as these columns do not contribute meaningful variability to the analysis. Subsequently, the missing values in integer columns are replaced with the mean of their respective columns, ensuring that the data remained representative of the overall dataset without introducing bias. This method of imputation is commonly used to retain valuable information while minimizing the potential impact of missing data. Finally, categorical variables are encoded to facilitate the machine learning process, transforming them into a numerical format that models can efficiently utilize.

### 3.3. Data Split into Genomic and Non-Genomic Sets

Genomic data refers to information derived from an organism’s DNA sequence, including mutations, chromosomal rearrangements, and gene expression profiles (8, 23, 29, 53). In contrast, non-genomic data encompasses a range of biological, clinical, or environmental factors, such as patient age, lifestyle factors, and medical history (26, 27, 38). In this step, we split the data into genomic and non-genomic categories that are suitable for machine learning models.

### 3.4. Feature Selection

In this step, feature selection is performed to identify the most important features contributing to model performance. We utilize a classifier to rank the importance of each feature, selecting the top features for both genomic and non-genomic datasets. The purpose of this step is to enhance the models’ efficiency by focusing on the most impactful features, which improves interpretability and reduces the risk of overfitting. By removing irrelevant or redundant features, we also decrease computational costs, ensuring that the model is both robust and effective for predicting PolyPhen and SIFT scores.

### 3.5. Correlation Analysis

To further refine the selected features, correlation analysis is conducted to identify and remove highly correlated features. This step is crucial in reducing multicollinearity, which can negatively impact model performance by causing overfitting and introducing redundancy in the feature set. By removing features that are strongly correlated, we aim to improve the model’s predictive power and generalizability. This refinement ensures that only the most relevant and distinct features are included, eliminating those that are highly correlated and could introduce multicollinearity.

### 3.6. Resampling

Resampling is used to adjust the distribution of input data types, typically to address class imbalances in classification problems (4, 14, 47). It involves modifying the dataset by either oversampling the minority class or undersampling the majority class, thereby ensuring a more balanced representation of classes within the data. By employing resampling methods, we prevent classification models from becoming biased toward the majority class, which is a common issue when dealing with imbalanced datasets.

### 3.7. Classification Models

The classification step is essential for the proposed approach because it significantly impacts the performance and accuracy of the classifiers in predicting PolyPhen and SIFT scores. In this study, we utilize three popular classification techniques: RF, XGBoost, and an ensemble model that combines RF and XGBoost. These techniques are well-suited for handling complex patterns and relationships within the genomic and non-genomic data, allowing the models to distinguish subtle differences between various outcomes. RF is known for its robustness and ability to manage high-dimensional data, while XGBoost is renowned for its efficiency and performance in handling imbalanced data. The ensemble model synergizes the strengths of both RF and XGBoost, leveraging their complementary capabilities to enhance predictive accuracy and generalization. By employing advanced classification techniques, we aim to investigate which dataset—genomic or non-genomic—provides better predictive performance for PolyPhen and SIFT scores.

### 3.8. Result Analysis

The analysis of the experimental results is to highlight the superiority of genomic data over non-genomic data in predicting PolyPhen and SIFT scores, emphasizing its relevance in mutation impact assessment. This analysis should demonstrate the potential of integrating genomic datasets with advanced machine learning techniques to improve the prediction accuracy of mutation pathogenicity, paving the way for enhanced research and clinical applications in disease detection and treatment.

## 4. EXPERIMENTAL SETUP

In this section, we present a comprehensive overview of the experimental framework employed in this research. We provide detailed information about the classification models used, along with the hyperparameters for each algorithm, ensuring transparency in the methodology. Additionally, we describe the genomic and non-genomic data, feature selection process, correlation analysis, and resampling techniques utilized in this study.

### 4.1. Classification Model Architecture

We implement three classification models, RF, XGBoost, and an ensemble RF-XGBoost model, to enhance predictive performance for PolyPhen and SIFT scores. Each algorithm is characterized by specific hyperparameters that significantly influence model effectiveness. The hyperparameters for each classification algorithm are summarized in **Table 1**, highlighting the strategic choices made to optimize the models’ performance. The RF model’s estimators and maximum depth allow for capturing complex relationships while maintain-ing a balance between bias and variance. In contrast, the XGBoost model’s estimators, learning rate, and maximum depth emphasize its ability to model intricate patterns in the data. For the ensemble RF-XGBoost model, to leverage the strengths of both models, we implement a Voting Classifier, which employs a soft voting strategy to combine predictions. In this approach, the predicted class probabilities from both the RF and XGBoost models are averaged, allowing the ensemble to consider the confidence of each model’s predictions. By using soft voting, the classifier considers the likelihood of each class, rather than just the predicted class label, thereby enhancing the robustness of the final decision.

**Table 1.** Classification Algorithms and Hyperparameters.

| Classification Algorithm | Hyperparameter | Value |
| --- | --- | --- |
| Random Forest (RF) | Number of Estimators | 1000 |
|  | Max Depth | 20 |
|  | Random State | 42 |
| Extreme Gradient Boosting Model (XGBoost) | Number of Estimators | 5000 |
|  | Learning Rate | 0.25 |
|  | Max Depth | 50 |
|  | Gamma | 0 |
|  | Sub Sample | 0.8 |
|  | Column Sample by Tree | 0.8 |
|  | Random State | 42 |
| Ensemble RF-XGBoost Model | Voting Strategy | Soft |
|  | Estimators | ['rf', 'xgb'] |

### 4.2. Genomic and Non-Genomic Data

We start with 556,268 rows and 186 columns from the TCGA data files, providing a wealth of information that require careful cleaning and transformation. After data cleaning, it results in a significant reduction of the dataset to 271,467 rows. After dropping the columns with a large number of missing values, it decreases the number of columns to 132. Columns that contained only one type of information are also eliminated as these columns do not contribute meaningful variability to the analysis, reducing the column count to 112. Here, 58 columns are genomic and 57 columns are non-genomic, of which PATIENT ID, PolyPhen, and SIFT columns are common in both datasets. It is important to note that PATIENT ID links the two datasets together. And, PolyPhen and SIFT are considered labeled features because they are the target we aim to predict. The missing values in integer columns are replaced with the mean of their respective columns, ensuring that the data remained representative of the overall dataset without introducing bias. Finally, categorical variables are encoded to facilitate the machine learning process, transforming them into a numerical format that models can efficiently utilize. **Table 2** and **Table 3** provide the names and descriptions of the genomic and non-genomic data, respectively.

**Table 2.** Description of Non-Genomic Data.

| No. | Non-Genomic Data | Description |
| --- | --- | --- |
| 1 | PATIENT_ID | Unique patient identifier |
| 2 | Hugo_Symbol | Gene name |
| 3 | Entrez_Gene_Id | Unique identifier for genes |
| 4 | Chromosome | Location of the gene on a chromosome |
| 5 | Start_Position | Starting position of the variant on the chromosome |
| 6 | End_Position | Ending position of the variant on the chromosome |
| 7 | Consequence | Biological effect of the variant |
| 8 | Variant_Classification | Type of variant (e.g., missense, nonsense) |
| 9 | Variant_Type | Type of genetic alteration (e.g., SNP, insertion) |
| 10 | Reference_Allele | Original allele at the variant position |
| 11 | Tumor_Seq_Allele1 | Allele from the tumor sequencing |
| 12 | Tumor_Seq_Allele2 | Second allele from the tumor sequencing |
| 13 | dbSNP_RS | Identifier for known single nucleotide polymorphisms |
| 14 | Matched_Norm_Sample_Barcode | Barcode identifier for matched normal tissue sample |
| 15 | Match_Norm_Seq_Allele1 | Allele from the matched normal sequencing |
| 16 | Match_Norm_Seq_Allele2 | Second allele from the matched normal sequencing |
| 17 | t_ref_count | Reference allele count in the tumor |
| 18 | t_alt_count | Alternative allele count in the tumor |
| 19 | n_ref_count | Reference allele count in the normal |
| 20 | n_alt_count | Alternative allele count in the normal |
| 21 | HGVSc | HGVS notation for coding sequence changes |
| 22 | HGVSp | HGVS notation for protein changes |
| 23 | HGVSp_Short | Short form of HGVS notation for protein changes |
| 24 | Transcript_ID | Identifier for the transcript affected by the variant |
| 25 | RefSeq | RefSeq identifier for this transcript |
| 26 | Protein_position | Position of the change in the protein |
| 27 | Codons | Codon changes caused by the variant |
| 28 | Allele | Specific alleles observed |
| 29 | Amino_acids | Changes in amino acids due to the variant |
| 30 | BIOTYPE | Biotype of transcript |
| 31 | CANONICAL | Indicates if the transcript is canonical |
| 32 | CCDS | Consensus coding sequence |
| 33 | CDS_position | Position in the coding sequence |
| 34 | CONTEXT | The reference allele per VCF specs |
| 35 | COSMIC | Overlapping COSMIC variants |
| 36 | DOMAINS | Protein domains affected by the variant |
| 37 | ENSP | Ensembl protein ID |
| 38 | EXON | Exon number where the variant is located |
| 39 | Existing_variation | Existing variation identifiers |
| 40 | Exon_Number | Number of the exon |
| 41 | Feature | Genetic feature affected by the variant |
| 42 | Gene | Name of the gene |
| 43 | HGNC_ID | HGNC identifier for the gene |
| 44 | IMPACT | Impact of the variant on the gene |
| 45 | NCALLERS | Number of callers that detected the variant |
| 46 | SOMATIC | Indicates if the variant is somatic |
| 47 | SWISSPROT | SwissProt identifier for the protein |
| 48 | SYMBOL | Gene symbol |
| 49 | FILTER | Filters applied to the variants |
| 50 | SYMBOL_SOURCE | Source of the gene symbol |
| 51 | TREMBL | TrEMBL identifier for the protein |
| 52 | UNIPARC | Universal protein resource cluster |
| 53 | all_effects | All effects predicted for the variant |
| 54 | cDNA_position | cDNA position of the variant |
| 55 | n_depth | Sequencing data coverage across a specific locus in a normal sample |
| 56 | t_depth | Sequencing data coverage across a specific locus in a tumor sample |
| 57 | PolyPhen | The predicted impact of a genetic variant on protein function |
| 58 | SIFT | The predicted impact of a genetic variant on protein function |

**Table 3.**
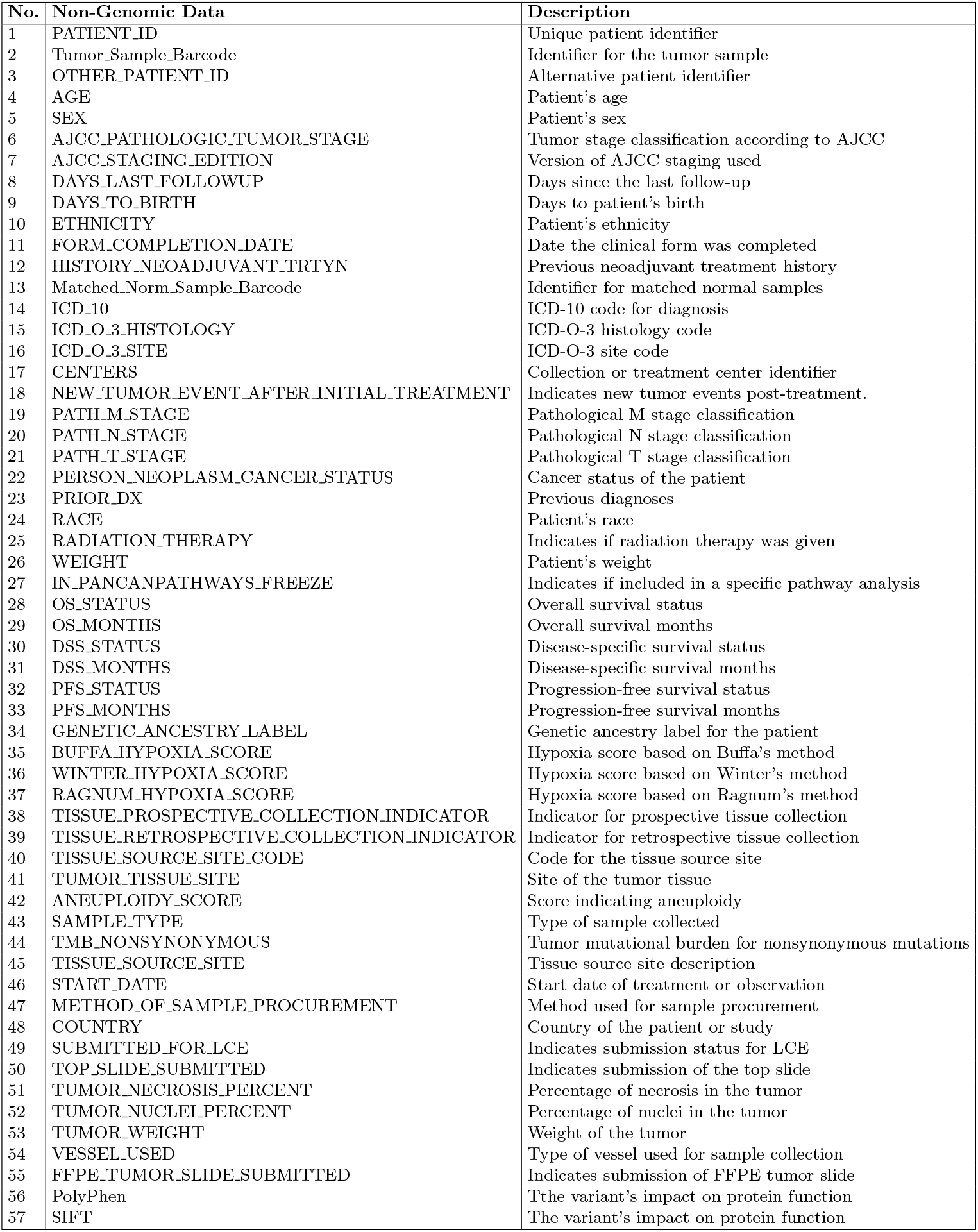
Description of Non-Genomic Data.

### 4.3. Feature Selection

**Table 4** shows the top 30 features selected by the RF classifier for both genomic and non-genomic data.

**Table 4.** Feature Selection for Genomic and Non-Genomic Data.

| Data Type | Top 30 Features |
| --- | --- |
| Genomic | PATIENT_ID, DOMAINS, HGVSp_Short, HGVSp, CONTEXT, Codons, cDNA_position, CDS_position, End_Position, Start_Position, Protein_position, Gene, UNIPARC, CCDS, HGNC_ID, Entrez_Gene_Id, HGVSc, all_effects, RefSeq, Exon_Number, EXON, t_ref_count, t_depth, SWISSPROT, Existing_variation, n_ref_count, n_depth, ENSP, Feature, and Transcript_ID |
| Non-Genomic | PATIENT_ID, dbSNP_RS, CENTERS, Tumor_Sample_Barcode, TMB_NONSYNONYMOUS, Matched_Norm_Sample_Barcode, OTHER_PATIENT_ID, WINTER_HYPOXIA_SCORE, TUMOR_WEIGHT, PFS_MONTHS, DAYS_TO_BIRTH, ANEUPLOIDY_SCORE, START_DATE, FORM_COMPLETION_DATE, AGE, BUFFA_HYPOXIA_SCORE, OS_MONTHS, DSS_MONTHS, RAGNUM_HYPOXIA_SCORE, SUBMITTED_FOR_LCE, DAYS_LAST_FOLLOWUP, PATH_T_STAGE, AJCC_PATHOLOGIC_TUMOR_STAGE, ICD_O_3_SITE, ICD_10, TUMOR_TISSUE_SITE, TUMOR_NUCLEI_PERCENT, WEIGHT, TUMOR_NECROSIS_PERCENT, and VESSEL_USED |

### 4.4. Correlation Analysis

Correlation analysis is conducted with a correlation threshold set at 0.70. **Figure 3** presents the feature correlation matrices for the genomic dataset before removing features. Features should be removed are: CDS position, Start Position, Protein position, EXON, t depth, SWISSPROT, n ref count, n depth, ENSP, Feature, and Transcript ID. After performing the correlation analysis, the genomic data features are reduced to 19 (from 30). By focusing on the final set of features, the classification models will have a clearer foundation to predict PolyPhen and SIFT scores, ultimately improving the model by reducing the risk of overfitting.

**Figure 3.**
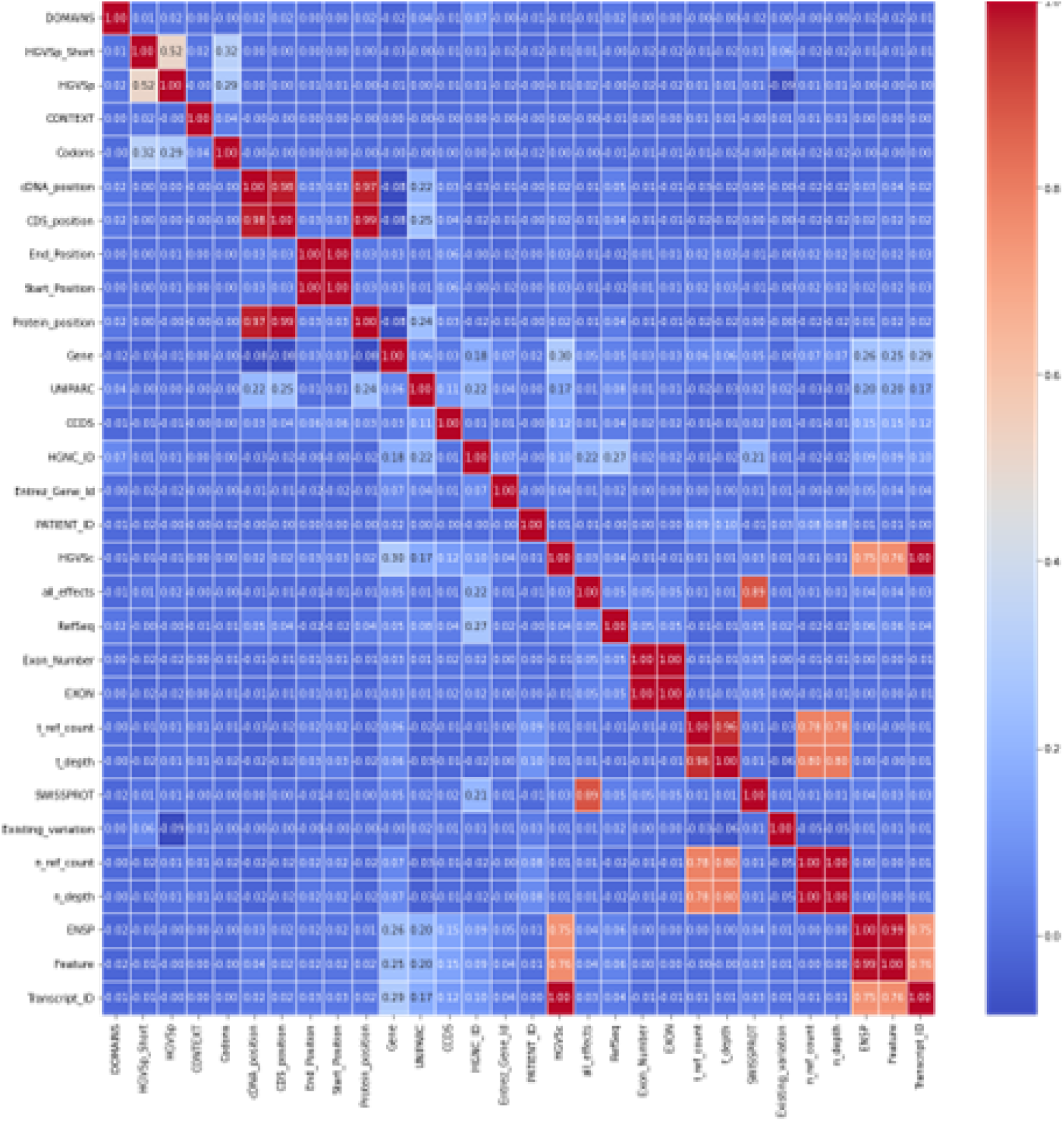
Genomic Correlation Matrix before Removal of Highly Correlated Features

**Figure 4**. presents the feature correlation matrices for the non-genomic dataset before removing features. Features should be removed are: Tumor Sample Barcode, Matched Norm Sample Barcode, START DATE, AGE, BUFFA HYPOXIA SCORE, OS MONTHS, DSS MONTHS, and ICD 10. After performing the correlation analysis, the non-genomic data features are reduced to 22 (from 30).

### 4.5. Resampling

In this research, we apply oversampling by duplicating the data points of the minority class to match the majority class. This approach enhances the model’s ability to learn from the minority class, ultimately improving its predictive performance. To illustrate the impact of resampling on our dataset, **Table 5** presents the distribution of data points before and after the resampling process for PolyPhen. This adjustment not only improves the overall balance of the PolyPhen dataset but also enhances the model’s ability to learn from and accurately predict instances from both classes.

**Table 5.** Distribution of Data Points Before and After Resampling for PolyPhen.

| Type | Benign | Probably Damaging | Possibly Damaging |
| --- | --- | --- | --- |
| Before Resampling | 114430 | 105187 | 51850 |
| After Resampling | 103700 | 103700 | 103700 |

**Figure 4.**
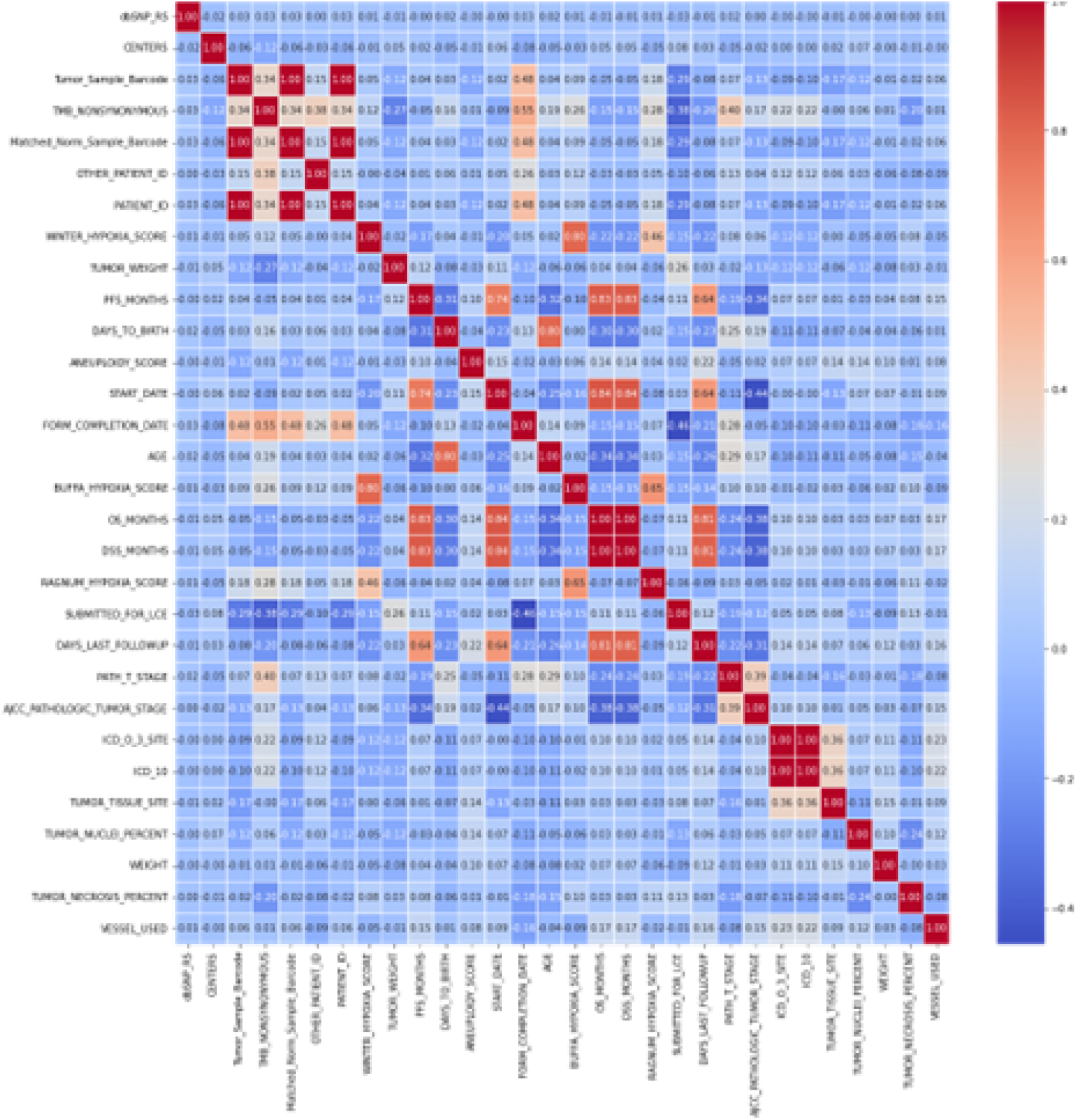
Non-Genomic Correlation Matrix before Removal of Highly Correlated Features

The SIFT dataset do not require resampling, as the number of samples is already adequate, which is detailed in **Table 6**.

**Table 6.** Data Distribution for SIFT (No Resampling Required)

| Type | Deleterious | Tolerated |
| --- | --- | --- |
| Data points | 114430 | 105187 |

## 5. RESULTS AND DISCUSSION

This section presents the experimental results obtained using both genomic and nongenomic datasets by applying three different machine learning models to predict the PolyPhen and SIFT scores. The results begin with the RF classifier, followed by the XGBoost and ensemble RF-XGBoost classifiers. For each of these models, we detail the accuracy and training time in the results. We aim to assess which dataset—genomic or non-genomic—yields superior predictive performance for PolyPhen and SIFT scores.

### 5.1. Performance of the RF Classifier

The performance of the RF classifier is evaluated for the dataset with 56 (excluding PolyPhen and SIFT) genomic and 55 (excluding PolyPhen and SIFT) non-genomic features after split, 30 genomic and 30 non-genomic features after feature selection, and 19 genomic and 22 non-genomic features after correlation analysis. **Table 7** highlights the metrics for PolyPhen and SIFT prediction using the RF classifier. Using 56 genomic and 55 non-genomic features for PolyPhen prediction, RF training time is 3590 seconds for genomic data and 1164 seconds for non-genomic data, with an accuracy of 81.93% for genomic data and 55.46% for non-genomic data. For SIFT prediction, the genomic data consistently outperforms the non-genomic data, achieving an accuracy of 89.03%, compared to 77.34% accuracy for non-genomic data. For both PolyPhen and SIFT, as the number of features decreases, there are minimal changes in accuracy, with decreases in training time. The results suggest that the smaller feature sets provide similar predictive power while being computationally more efficient.

**Table 7.**
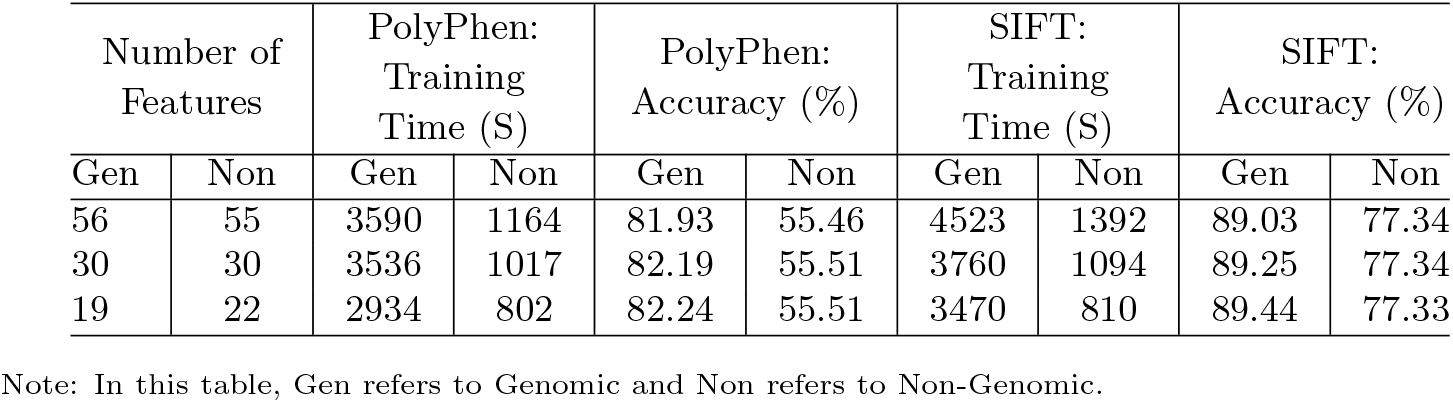
Metrics for PolyPhen and SIFT Prediction using RF Classifier.

### 5.2. Performance of the XGBoost Classifier

The performance of the XGBoost classifier is assessed similarly to that of the RF classifier. **Table 8** highlights the metrics for PolyPhen and SIFT prediction using the XGBoost classifier. Utilizing 30 genomic and 30 non-genomic features for PolyPhen prediction, XGBoost training time is 1569 seconds for genomic data and 3836 seconds for non-genomic data, achieving an accuracy of 88.21% for genomic and 55.36% for non-genomic datasets. For SIFT prediction, the genomic data significantly outperforms the non-genomic data, attaining an accuracy of 95.03% compared to 77.23% accuracy for non-genomic data. Like RF, the results indicate that the smaller number of features maintains similar predictive power while enhancing computational efficiency.

**Table 8.** Metrics for PolyPhen and SIFT Prediction using XGBoost Classifier.

| Number of Features |  | PolyPhen: Training Time (S) |  | PolyPhen: Accuracy (%) |  | SIFT: Training Time (S) |  | SIFT: Accuracy (%) |  |
| --- | --- | --- | --- | --- | --- | --- | --- | --- | --- |
|  |  | Gen | Non | Gen | Non | Gen | Non | Gen | Non |
| 56 | 55 | 1921 | 5309 | 88.33 | 55.35 | 1592 | 2913 | 95.06 | 77.23 |
| 30 | 30 | 1569 | 3836 | 88.21 | 55.36 | 967 | 2596 | 95.03 | 77.23 |
| 19 | 22 | 1217 | 3372 | 88.12 | 55.39 | 727 | 1983 | 95.04 | 77.25 |

Note: In this table, Gen refers to Genomic and Non refers to Non-Genomic.

### 5.3. Performance of the Ensemble RF-XGBoost Classifier

**Table 9** summarizes the experimental results for PolyPhen and SIFT prediction using the ensemble RF-XGBoost model. Using 19 genomic and 22 non-genomic features for PolyPhen prediction, the ensemble training time is 2878 and 6382 seconds, respectively. The accuracy achieved is 88.30% for genomic and 55.49% for non-genomic data. For SIFT prediction, genomic data consistently outperforms non-genomic data, achieving an accuracy of 95.12%, compared to 77.32% for non-genomic data. The ensemble RF-XGBoost model delivers the best performance in predicting both PolyPhen and SIFT scores, achieving the highest accuracy across both genomic and non-genomic datasets. However, the ensemble model requires longer training time compared to the individual models (e.g., RF and XGBoost).

**Table 9.** Metrics for PolyPhen and SIFT Prediction using RF and XGBoost Classifier.

| Number of Features |  | PolyPhen: Training Time (S) |  | PolyPhen: Accuracy (%) |  | SIFT: Training Time (S) |  | SIFT: Accuracy (%) |  |
| --- | --- | --- | --- | --- | --- | --- | --- | --- | --- |
| Gen | Non | Gen | Non | Gen | Non | Gen | Non | Gen | Non |
| 56 | 55 | 5632 | 9642 | 88.43 | 55.45 | 5166 | 3977 | 95.13 | 77.31 |
| 30 | 30 | 4514 | 323 | 88.42 | 55.47 | 4966 | 3778 | 95.14 | 77.31 |
| 19 | 22 | 2878 | 6382 | 88.30 | 55.49 | 4033 | 2829 | 95.12 | 77.32 |

Note: In this table, Gen refers to Genomic and Non refers to Non-Genomic.

### 5.4. Performance Analysis

After applying feature selection, correlation analysis, and resampling, the results for RF, XGBoost, and ensemble RG-XGBoost models using genomic and non-genomic datasets are shown in **Table 10**. According to the simulation results, the XGBoost model performs significantly better on genomic datasets compared to non-genomic datasets across accuracy and training time. On the genomic dataset, XGBoost achieves an accuracy of 88.12% for PolyPhen with a training time of 1,217 seconds, and 95.04% for SIFT with a training time of 727 seconds. On the non-genomic dataset, it requires 3,372 seconds for PolyPhen and 1,983 seconds for SIFT, achieving accuracies of 55.39% and 77.25%, respectively. While the ensemble RF-XGBoost model shows comparable accuracy to XGBoost, it requires significantly more training time without offering a notable improvement in performance.

**Table 10.** Metrics for PolyPhen and SIFT Prediction using Data after Resampling.

| Classification Model | PolyPhen or SIFT | Genomic Dataset |  | Non-Genomic Dataset |  |
| --- | --- | --- | --- | --- | --- |
|  |  | Training Time (S) | Accuracy (%) | Training Time (S) | Accuracy (%) |
| RF | PolyPhen | 2934 | 82.24 | 802 | 55.51 |
|  | SIFT | 3470 | 89.44 | 810 | 77.33 |
| XGBoost | PolyPhen | 1217 | 88.12 | 3372 | 55.39 |
|  | SIFT | 727 | 95.04 | 1983 | 77.25 |
| Ensemble RF-XGBoost | PolyPhen | 2878 | 88.30 | 6382 | 55.49 |
|  | SIFT | 4033 | 95.12 | 2829 | 77.32 |

Detailed comparisons of training time and accuracy due to genomic and non-genomic datasets using classification models, specifically for classifying PolyPhen and SIFT scores, are not well-documented in the published literature (B. Haque et al., 2025; S. Agajanian et al., 2019; S. Das et al., 2025; K. Thedinga and R. Herwig, 2021). Our study reveals that, based solely on accuracy, genomic datasets are the best choice for all three classification models while classifying PolyPhen and SIFT scores. However, when considering both accuracy and training time, genomic datasets are the most favorable option only for the XGBoost model. For the RF and ensemble RF-XGBoost models, predictions for both PolyPhen and SIFT scores yield higher accuracy but require significantly more training time.

## 6. CONCLUSIONS

Recent studies suggest that there are no reliable ready-to-use genomic datasets to explore the effectiveness of machine learning models to accurately identify potentially harmful mutations and support research or clinical decision-making. This research presents a method to construct and assess the impact of cancer genomic datasets for PolyPhen and SIFT classifications using ensemble learning. We use the TCGA datasets from cBioPortal and popular RF, XGBoost, and ensemble RF-XGBoost models. Through a systematic process of data preprocessing, feature selection, correlation analysis, and resampling, we refine the genomic and non-genomic datasets to improve model generalizability and reduce computational costs. The experimental results confirm that the genomic data consistently outperforms the non-genomic data across all models and features, with the ensemble model demonstrating the highest predictive accuracy for both PolyPhen and SIFT classification. The findings indicate that the ensemble RF-XGBoost model performs best when applied to genomic data, achieving average accuracies of 88.43% for PolyPhen and 95.13% for SIFT. It should be noted that the performance gain comes at the cost of increased training time, highlighting a key trade-off between accuracy and computational efficiency. This research provides valuable insights into the application of machine learning models for predicting PolyPhen and SIFT scores, which can contribute to advancements in genomic research and medical interventions. In our next endeavor, we intend to examine a wider array of genomic datasets from TCGA and other sources to enhance the robustness of prediction models across diverse genetic backgrounds.

## SUMMARY POINTS

1. This research constructs genomic and non-genomic datasets from TCGA to assess the impact of genetic mutations in cancer treatment.
2. PolyPhen and SIFT scores are used to assess how amino acid substitutions affect protein function and mutation pathogenicity.
3. Machine learning classifiers—RF, XGBoost, and an ensemble RF-XGBoost model—are evaluated for improved prediction accuracy.
4. Genomic data outperforms non-genomic data, with the RF-XGBoost ensemble achieving the highest accuracy for PolyPhen and SIFT scores.
5. The research highlights the potential of artificial intelligence in genetic data analysis for treating diseases such as cancer.

### FUTURE ISSUES

1. Focus on incorporating more diverse genomic datasets from different populations to improve the generalizability and robustness of prediction models across various genetic backgrounds.
2. Develop clinically applicable tools that integrate machine learning models into healthcare systems, concerning data privacy, processing speed, and interpretability for medical professionals.
3. Combining genomic data with other omics data (e.g., transcriptomics, proteomics, and metabolomics) could enhance prediction accuracy and provide a more comprehensive understanding of mutation effects on disease mechanisms.

## ACKNOWLEDGMENTS

The authors express gratitude to the Computer Architecture and Parallel Programming Laboratory (CAPPLab) in the College of Engineering at Wichita State University for their support in conducting machine learning simulations on The Cancer Genome Atlas (TCGA) datasets. The authors are extremely grateful to the National Institutes of Health (grant no. 1R01CA260958-01A1 to M.J.U.) and the Phi Beta Psi Sorority Trust (grant nos. AWD00000652 and AWD00001248 to M.J.U.) for partially supporting this study.

